# Evolutionary and Deep Learning Models Highlight Deleterious Mutations Behind the History of Sugar Beet Breeding

**DOI:** 10.64898/2026.03.18.712733

**Authors:** Evan Long, Rajtilak Majumdar, Rachel Naegele, Kevin Dorn

## Abstract

Sugar beet (*Beta vulgaris ssp. vulgaris*) is a major global source of sucrose. Domestication and a history of breeding have shaped the distribution of deleterious mutations across the genome. Accurately resolving these variants offers a compelling avenue to accelerate crop improvement through breeding or genetic engineering. We combine three scales of evolutionary time: protein conservation defined by conservation across all living organisms (SIFT), angiosperm-wide DNA language deep learning model (PlantCaduceus), and fine-scale evolutionary rates from Amaranthaceae plant family sequence alignments to define a high-confidence set of deleterious variants enriched for alleles under negative selection. We then evaluate these deleterious mutations across a multi-study panel of more than 1,900 whole-genome sequenced wild and cultivated beet accessions. We found that domesticated beet accessions exhibit higher deleterious load relative to wild maritima, yet sugar beet carries significantly fewer deleterious variants than other cultivated beet crops. Historical trends show a decline in genetic load across more than a century of sugar beet entries in the U.S. NPGS, reflecting sustained purging during modern breeding. This combinatorial method of leveraging evolutionary conservation reveals how deleterious mutations have shaped sugar beet breeding history and provides potential targets for future selection, causal-variant discovery, and precision genome engineering.

## Introduction

*Beta vulgaris ssp. vulgaris* refers to a species that encompasses several cultivated beet crops, including table beet, fodder beet, chard, and sugar beet. Sugar beet is a major source of sucrose and an important component of agricultural systems in temperate regions (Asadi 2006). As demand for sustainable yield gains intensifies under evolving production constraints, breeding programs increasingly leverage genomic tools to accelerate genetic improvement while preserving agronomically important diversity (Monteiro *et al*. 2018; Subrahmanyeswari and Gantait 2022). Sugar beet is predominantly outcrossing, and modern breeding widely exploits the performance advantages of heterozygous hybrids (Hallahan *et al*. 2018). This mating system has implications for the evolutionary fate of deleterious mutations, in the context of both natural and breeding history.

The history of beets (*Beta vulgaris*) goes back more than 2000 years. The wild ancestors to modern beets (*Beta vulgaris ssp. maritima*) were neither spherical nor succulent. Early depiction by Roman Naturalist Pliny indicated sea beets as less fleshy than saffron crocus, hinting some swelling of roots (Dalby 2003). Earlier reports indicate that beet was predominantly a leaf crop and modern-day Swiss Chard represents such leafy and domesticated forms of *B. vulgaris*. Domestication of *B. vulgaris* is believed to have focused on selection that triggered the expansion of cambial layers (Goldman and Janick 2021). Thickened and succulent roots for food may have appeared around the mid-1500s in Italy or Germany. Selection of modern-day more tender, spherical, and sweeter roots continued over several centuries and became highly prevalent in Europe during in the 19th century. Fodder beets, then known as mangel-wurzel or mangolds, believed to have originated in Germany or the Netherlands in the mid-1500s Though, large-scale cultivation started sometime in the late 1700s (Melzer *et al*. 2013). Sugar beet underwent a more recent bottleneck (Strong selection on sugar) in the mid-1700s. Discovery of sugar beet (then known as “Runkelrüben”; cane sugar) by A.S. Marggraf in 1747 as a plant species that stores sucrose in the roots, incentivized growers to increase production of this crop. Sucrose content (fresh weight basis) in the roots was ∼4% during 1700s. Later in early 1800s, selection for root biomass and sucrose content occurred mainly through mass selection rather than progeny testing that came later. This resulted in significant increase in root biomass and sucrose content (∼17-19%) in commercial sugar beet varieties.

Continual breeding and selection often confer benefits in part through the indirect purging of genetic load, as improvement efforts select against deleterious mutations that reduce fitness or performance (Yang *et al*. 2017; Charlesworth 2018). In outcrossing species, recessive deleterious variants are often masked by heterozygosity, which can mitigate their phenotypic effects in hybrids while contributing to heterosis through complementation (Sun *et al*. 2023). However, the same masking can hinder purging of recessive load, and periods of domestication or directional selection may further increase the frequency of linked deleterious alleles via hitchhiking (Hartfield and Otto 2011) or through reductions in effective population size (Gaut *et al*. 2018). One result of these deleterious mutations is slow progress in breeding gains due to inadvertent selection of their complementation rather than removal.

To manage genetic load, breeders have long used strategies such as selection among inbred lines and hybrid breeding (Dwivedi 2023). The breeding of inbred lines allows for recessive deleterious mutations to be exposed to selection, while the generation of a final hybrid leverages complementation to achieve optimal performance. In practice, strong inbreeding depression, complex trait architectures, and the population specificity of quantitative trait loci (QTL) can limit the effectiveness of pedigree or marker-assisted approaches. Prioritizing the removal of deleterious variants via breeding and genetic engineering could provide a method to accelerate genetic gains, while reducing the need to evaluate phenotypes in field environments for every generation. Previous studies have shown that the incorporation of functional information such as deleterious mutations can improve genomic prediction of fitness related traits, especially in unrelated germplasm (Long *et al*. 2023; Wu *et al*. 2023). This is especially important as one avenue to tackle modern abiotic and biotic stresses is to introduce novel traits from wild relatives (Monteiro *et al*. 2018; Mulet 2022), likely also introducing more deleterious mutations into productive varieties.

Evolutionary conservation has proven useful for predicting deleteriousness in a population agnostic framework: sites conserved across species are more likely to be functionally constrained, and derived variants at these sites are more likely to be deleterious. Functional impact on protein structure can also be used to augment functional predictions within coding regions of the genome (Long and Monroe 2025). Recently, deep learning models, such as PlantCaduceus (Zhai *et al*. 2025b) or Evo2 (Jiang *et al*. 2025), have been built on evolutionary information to generate models capable of variant effect prediction. This genre of models has been shown to aid in the discovery of causative mutations in maize (Wang *et al*. 2025).

In this study, we identify putative deleterious variants in *Beta vulgaris* using evolutionary and functional annotations, evaluate the distribution of these deleterious variants across cultivated and wild beets, and explore their putative role in historic breeding efforts in sugar beet. Understanding the nature of these deleterious mutations can offer future targets for genetic improvement and help accelerate the purging of genetic load during crosses and introgression from wild germplasm.

## Methods

### Variant Calling

Raw reads for 1910 beet and related species were downloaded from National Center for Biotechnology Information (NCBI) sequence read archive (see Table S1 for accession IDs, Table S2: Species). Metadata including subspecies crop type (i.e. chard, fodder beet, table beet, sugar beet), and location of origin were extracted from corresponding publication metadata or from the National Plant Germplasm System (NPGS) and Leibniz Institute of Plant Genetics and Crop Plant Research (IPK) databases. These sequences are derived from six different sequence projects with varied germplasm sources, sampling methods, sequencing depths, and study goals (Table S3) (Galewski and McGrath 2020; Wang *et al*. 2023; Wang *et al*. 2024; Dhiman *et al*. 2025; Reeves *et al*. 2026). Sequence reads were trimmed using trimmomatic, (Bolger *et al*. 2014) and aligned to the EL10.2 *Beta vulgaris* reference genome (McGrath *et al*. 2023) using bwa-mem (Li 2013). Using samtools V1.20 (Li *et al*. 2009), alignments were processed using fixmate and markdup, then indexed. Variants were then called using DeepVariant (Yun *et al*. 2020) version 1.8.0 (WGS setting). Individual gvcf files were then combined using GLnexus (configuration “DeepVariantWGS”).

To allow for varied data sources, while maintaining high confidence in variant calls, filtering and imputation were performed. Genotypes calls with <5 reads supporting the call or a genotype quality score <20 were removed. Sites were then filtered for a minimum of 1% minor allele frequency and minimum proportion of accessions with genotypes calls at a site of 10%. Genotypes were then imputed using BEAGLE (Browning *et al*. 2018) under default parameters.

### Evolutionary Conservation

Genomes were gathered from across the Amaranthaceae (sometimes classified as Chenopodiaceae) plant family including 85 assemblies from over 70 different species (Table S4), using the NCBI datasets tool (https://www.ncbi.nlm.nih.gov/datasets/genome/). Genes were aligned via multiple sequence alignment using a previously developed pipeline https://bitbucket.org/bucklerlab/p_reelgene/ (Long *et al*. 2023; Schulz *et al*. 2025). Briefly, EL10.2 reference transcripts (https://phytozome-next.jgi.doe.gov/) were aligned to each genome to fetch corresponding homologous exonic regions. Exonic regions whose alignment length was at least 90% of the total gene length of each transcript and had the highest alignment score were retained for multiple sequence alignment. Genic regions were used for multiples sequence alignment to enable proper alignment across evolutionary time, as multiple sequenece alginment becomes increasingly difficult at this evolutionary scale as the distance from genes increases. MAFFT (Katoh and Standley 2013) was used to generate multiple sequence alignments for each EL10.2 reference transcript (--ep 0 --genafpair --maxiterate 1000). Gapped positions in the reference transcript were removed to simplify in-frame protein coding analysis. Gene trees were then using RAxML (Stamatakis 2014), and calculated evolutionary rates using baseml from the PAML (Yang 2007) suite of tools. This provided an evolutionary rate for each base in each transcript in the Beet genome.

### Deleterious Annotation

A combination of tools and metrics were used to identify deleterious mutations. A Sorting-Intolerant-From-Tolerant (SIFT) mutation database was generated for EL10.2 https://github.com/rvaser/sift4g (Vaser *et al*. 2016), using the UniRef90 protein database. Additionally, we annotated our variants using the PlantCAD2 (Zhai *et al*. 2025b) DNA foundation model for angiosperms (https://github.com/plantcad/plantcad, model: PlantCAD2-Large). Finally, for a conservative method of determining a selection of deleterious mutations, we retained single nucleotide polymorphisms (SNPs) with a SIFT score ≤0.05, an evolutionary rate ≤0.5, and a PlantCAD2 zero-shot score in the bottom one percentile for both Single nucleotide polymorphisms (SNP) and indels.

### Data Correlations

The abundance of each classified deleterious mutation was calculated, and a genetic load was calculated for all mutations, heterozygous mutations, and homozygous mutations for each accession. Linear regression was performed between genetic load and sugar beet accessions with entry years recorded in the U.S. NPGS, and Pearson correlations are reported. Pairwise t-tests between genetic load abundance among *Beta vulgaris* subspecies and crop types were performed. For studies where biomass and sugar were available (Wang *et al*. 2024), trait and genetic load Pearson correlations were calculated and displayed using the R package *corrplot* (Wei *et al*. 2017).

## Results

### Panel of Sugar Beet Accessions

We gathered sequence reads from over 1900 *Beta vulgaris* accessions, including a few related species, and processed them in using a unified QC, read mapping, and variant calling pipeline. These accessions were sequenced as part of six different sequencing projects (Table S3) and are derived from cultivated and wild samples from all over the world (Fig. 1). The resulting genotype files containing all accessions included over 38M variants, however these were filtered down to a higher confidence set of 9M variants, of which ∼480k fell into gene coding regions of the genome that were for evaluated for predicted deleterious effects.

**Figure 1.**
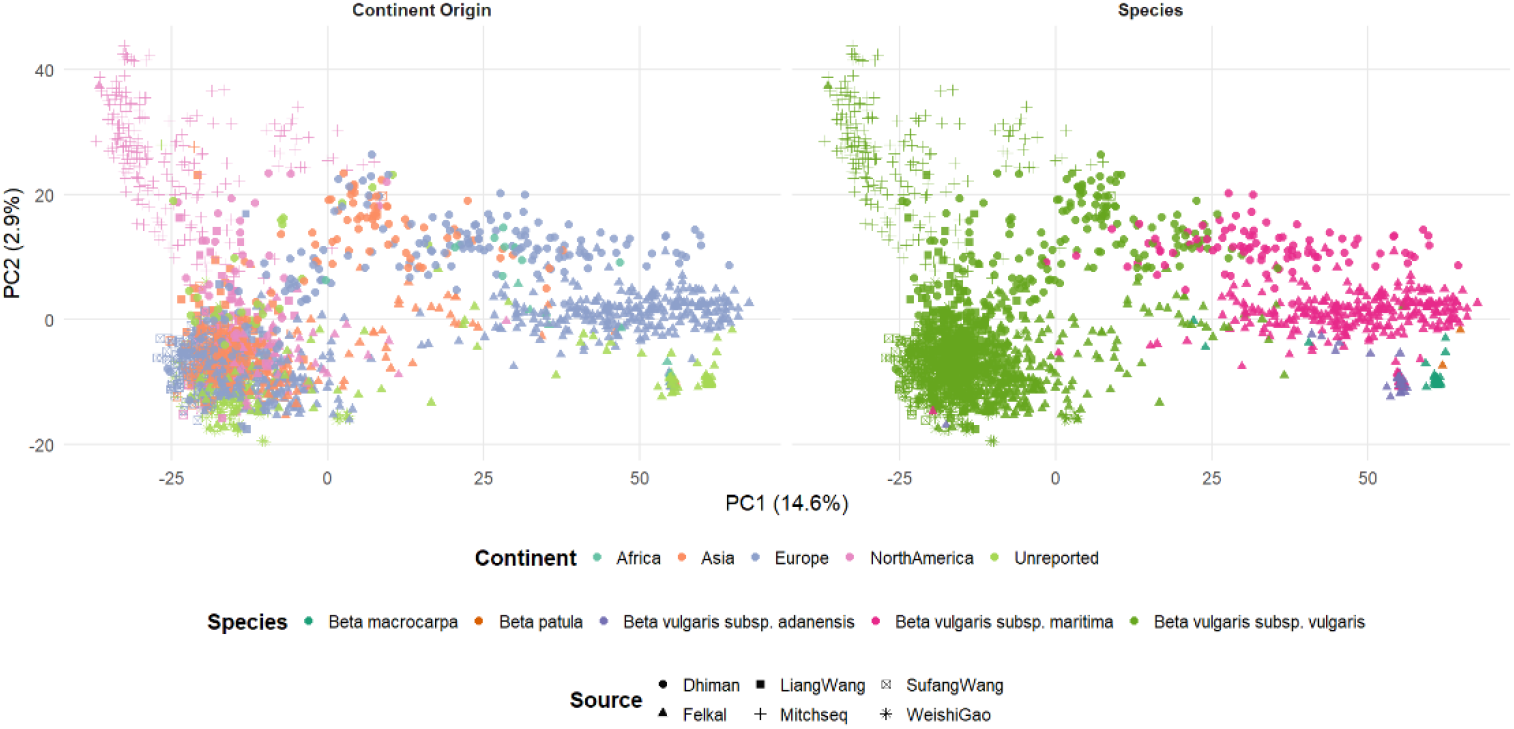
Principal Component Analysis of Beet Panel. Principal component analysis of genotype data. Both plots display the same accessions: colors display the continent from which the sample originated (left) and the species to which each sample belong (right). The genotyping study from which each genotype is derived is shown as the point shape. The percentage of variation explained by each principal component is shown in parentheses.

### Deleterious Annotations

We utilized three methodologies to identify putative deleterious mutations across the beet genome among over 1900 accessions. Mutations flagged as deleterious by SIFT showed evidence of negative selection across the large panel of beet accessions (Fig. S1). Those deleterious mutations classified by PlantCAD2 also showed evidence of strong negative selection (Fig. S2). These results showed consistency with evolutionary conserved sites (Figs. S1-2). We found an overlap of 1,926 deleterious mutations classified by all three methods (Fig. 2A), that exhibited strong enrichment for rare allele frequencies (Fig. 2B). Aside from a few potential hot spots, the distribution of the deleterious mutations largely matched the genome-wide distribution of genic variants (Fig. S3).

**Figure 2.**
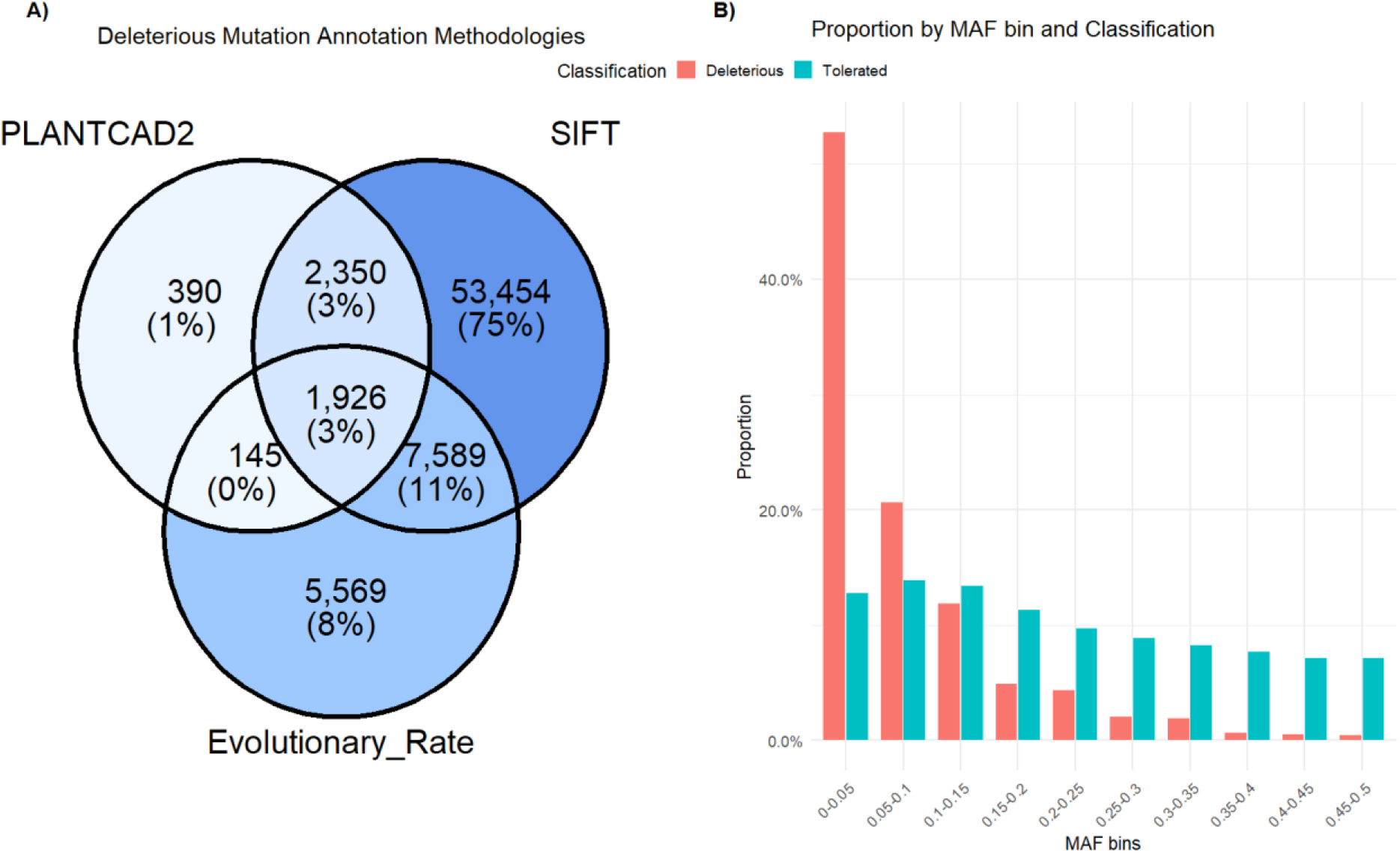
Deleterious Annotations. A) The number of mutations classified by each of the three evolutionary methods PlantCadaceus (PLANTCAD2), Sorting-Tolerant-From-Intolerant (SIFT), and PAML (Evolutionary Rate). The center portion represents the ∼2k deleterious mutation sites used in this study where all three methods predicted a deleterious effect. B) The folded site frequency spectra for the classified deleterious and tolerated mutations. The proportion of mutations with the corresponding minor allele frequency (MAF) within each classification are displayed.

### Deleterious Mutations Accumulation In *Beta vulgaris*

The abundance of these deleterious mutations, classified by all three of the used annotation methods, shows distinct variability across the *Beta vulgaris* accessions (Fig. 3). Within the *Beta vulgaris* species, the total number of deleterious mutations in cultivated beets, *vulgaris*, and the wild sea beets, *maritima*, was significantly higher than the wild *adanensis* population (Fig. 3A). However, the number of homozygous mutations exhibited a nearly opposite trend with the cultivated beets having significantly lower accumulation, and the *maritima* population having less than the *adensis* population. And the converse is true for heterozygous mutations with the cultivated beets containing the highest amount of heterozygous genetic load. Within the cultivated beet populations, sugar beets manifest significantly fewer deleterious mutations than table beets, fodder beets, and chard (Fig. 3B).

**Figure 3.**
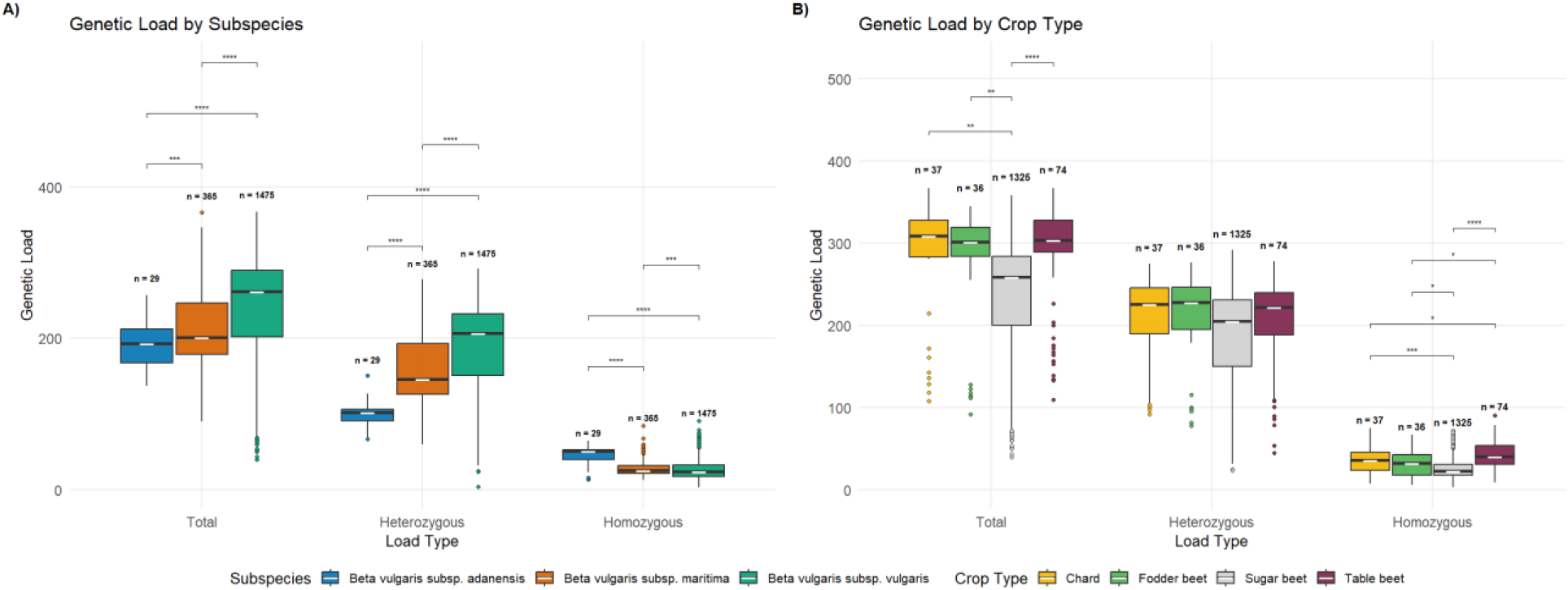
*Beta vulgaris* Deleterious Mutations. A) The genetic load (number of deleterious mutations) for each accession belonging to each subspecies within *Beta vulgaris* are displayed. Genetic load is broken into three different classifications 1-the total number of deleterious mutations, 2-the number of heterozygous deleterious mutations, 3-the number of homozygous deleterious mutations. B) The genetic load of each crop type (within *Beta vulgaris ssp vulgaris*) is displayed. Adjusted p-values significance levels are shown: * ≤ 0.05; **≤ 0.01; ***≤ 0.001; ****≤ 0.0001

### Genetic Load and Sugar Beet Breeding

Many of the sugar beets sequenced in this dataset are stored in the U.S. NPGS along with metadata corresponding to their acquisition. Using the sample entry date, we attempted to evaluate how genetic load may vary throughout the breeding history of sugar beet. We found that there are significant negative correlations between our assessment of genetic load and the year the variety was submitted to the germplasm system (Fig. 4). This correlation is significant for both the heterozygous mutations and, while diminished, homozygous mutations.

**Figure 4.**
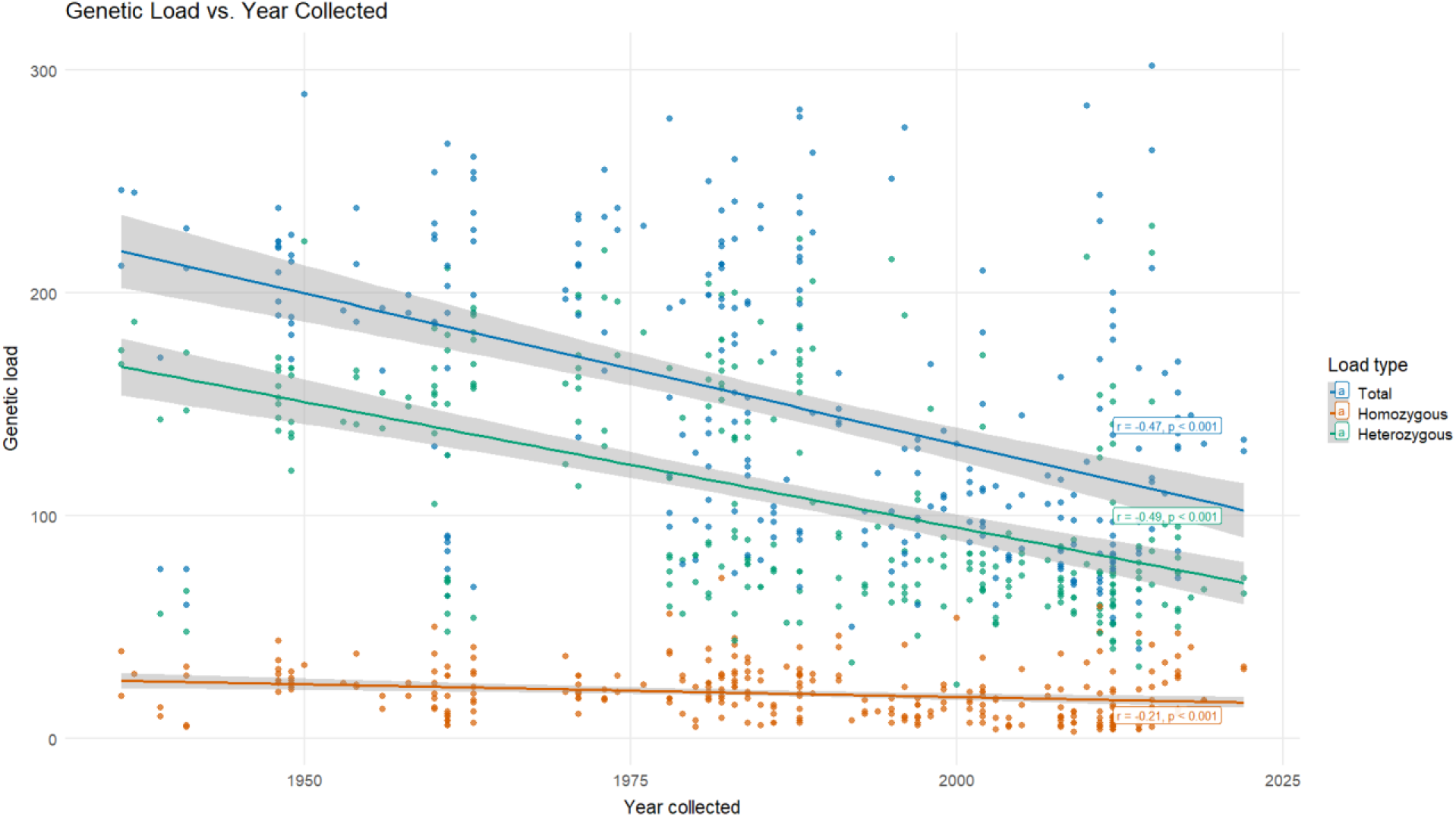
Deleterious Mutations in Sugar Beet by Collection Year in the National Plant Germplasm System. The genetic load (number of deleterious mutations) for each sugar beet accession with accompanying collection dates are displayed. Genetic load is broken into three different classifications 1-the total number of deleterious mutations, 2-the number of heterozygous deleterious mutations, 3-the number of homozygous deleterious mutations. Linear regressions, correlation coefficients (r), p-values, and corresponding 95% confidence intervals (shaded regions) are shown.

While the amount of genotypic data in this sequenced panel of *Beta vulgaris* is large, the amount of trait related data is limited, especially for fitness related traits where genome-wide genetic load has expected impacts. One subset of the sugar beet data originating from a study by Sufang Wang et al. (Wang *et al*. 2024) does contain many phenotypic observations including total biomass, sugar content, and related root traits (Fig. 5). We found a small, but significant negative correlation between homozygous load and total growth-related traits such as biomass, compactness (ratio of maximum root diameter to root length, and top projection area (an estimate of total root surface area based on multi-view images of tap roots). Genetic load also showed some positive correlations with root sugar content.

**Figure 5.**
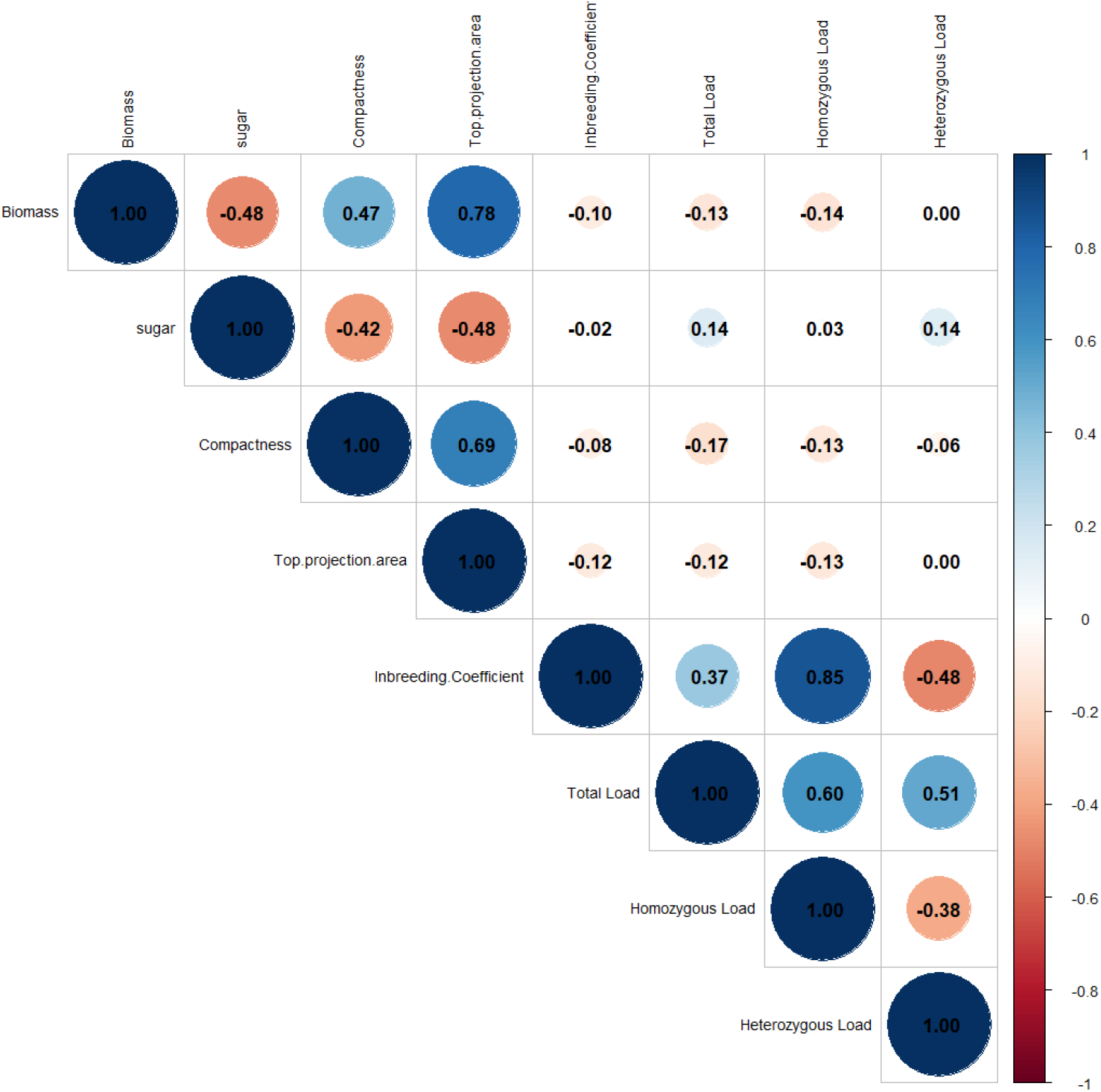
Correlations Between Traits and Genetic Load. Pearson correlations between traits and genetic load for a subset of ∼200 sugar beet accessions are displayed. Circle size and color correspond to the magnitude and direction of the correlation. Trait ontologies: biomass-total measured mass, sugar-percent sugar content, compactness-ratio of maximum root diameter to root length, top projection area-an estimate of total root surface area based on multi-view images of tap roots.

## Discussion

### Genetic Diversity and Data Integration

This study pulled whole genome sequence data from multiple sources in an effort to leverage one large dataset representing *Beta vulgaris* genetic diversity. While sources for this sequence data varied, a common genotyping pipeline was used to minimize biases that would have been introduced by trying to merge called genotypes from each individual study. The robust size of this genotype panel enables stronger statistical power to detect important variation.

Across the diversity of the accessions, the distinction between the *Beta vulgaris* subspecies, as well as the closer relationship between the wild progenitor *Beta vulgaris ssp. maritima* and modern cultivated beets can be observed compared to *Beta vulgaris ssp. adanensis*, consistent with earlier studies (Fig. 1, PC1 14.6% variation explained) (Tehseen *et al*. 2024). While to a lesser extent, the second axis of variation (PC2, 2.9% variation explained) seems to be driven by the sequencing study from which the accession was derived. This is visible in both the *maritima* population division between Felkal et al and Dhiman et al., as well as in the cultivated beets with the *Mitchseq* data separated from the other cultivated accessions. Between *maritima* populations, the separation may be due to Dhiman et al. samples having been sequenced at a higher depth (∼15X vs ∼6X; Table S3) compared to Felkal et al. Among cultivated lines, the separation may correspond to the pooled sampling method and/or higher sequencing depth of the *Mitchseq* data. It is important to note that many sugar beet database entries in public databases exhibit genetic heterogeneity within accessions due to their heterozygous, outcrossing nature, so even samples of the same accession are not likely to have identical genotypes (Galewski and McGrath 2020). Another explanation could be divergent selection pressure among the accessions that are primarily USDA derived pre-breeding material selection as sources of novel disease resistance. These results highlight the difficulty of integrating data from various sources, as there likely does not exist any specific data filters and imputation methodology to remove all biases, or an effort to do so may in fact remove biologically meaningful signal.

### Deleterious Annotation Methods

Multiple methodologies were leveraged to generate a strong classification of deleterious mutations. SIFT is a commonly used tool in measuring protein coding mutations, relying on evolutionary conservation of amino acid sequences from a large span of organisms (Vaser *et al*. 2016). Similarly, PlantCAD2 relies on evolutionary information, however it leverages a deep learning, DNA language model with single-nucleotide resolution trained on 65 angiosperm genomes (Zhai *et al*. 2025a). This DNA language model utilizes 8192bp of context around the mutation to assign mutation a “zero-shot score” corresponding to its difference in log-likelihoods of the alleles, with a high negative values indicating strong evolutionary conservation. Finally, the evolutionary rate generated from gene-level multiple sequence alignments provides the power and context of much more related species than the other two methods. Each method utilizes a different scale of evolutionary time to the detection of deleterious mutations: Protein database of all living things (SIFT) -> Angiosperm plants (PlantCAD2) -> Amaranthaceae family (multiple sequence alignment). Our combined method produced a conservative set of the most likely deleterious mutations we could detect. The deleterious nature of these mutations is supported by the measurable negative selection across the population (Fig. 2B).

The advent of deep learning methodologies provides strong resolution in the realm of variant effect prediction. The substantially greater number of unique variants classified by SIFT as deleterious suggests a possible tendency towards false‐positive calls when relying solely on distant evolutionary conservation (Fig. 2A). The power that is achieved by the deep learning and deep evolutionary time of the PlantCAD2 method is highlighted by its large overlap with the other two methods. The threshold for which we classified PlantCAD2 scores as deleterious was chosen for a strong enrichment, however the depth of this model allows for easier ranking of variant effects, where SIFT typically follows a strong binary distribution (Fig. S4).

For this study we restricted ourselves to the evaluation of deleterious mutations in gene coding regions. This restriction is due to the secondary support for mutation effects given by predictable effects on proteins. The underlying principle behind these methods is also a multiple sequence alignment, which is much less reliable outside of gene coding regions of the genome. In the coming years deep learning algorithms may excel to a point where intergenic assessments of mutations effects are accurate and reliable, but we made no attempt to test those methods in this study.

### Genetic Load In Beets: Past, Present, and Future

The measurement of the accumulation of these deleterious mutations across this large panel of diverse accessions reveals the effects of both domestication and recent breeding history. The work presented here highlights evolutionary adjustments of deleterious variants in the genome of modern-day sugar beet accessions and is in line with the cost-of-domestication hypothesis (Moyers *et al*. 2018). Our results highlight that domestication resulted in increased genetic load in culitivated beet lines compared to their wild counterparts (Fig. 3A), observations that have been reported in other naturally outcrossing crops such as maize (Lozano *et al*. 2021) and cassava (Ramu *et al*. 2017). Most of this disparity in genetic load can be attributed to heterozygous mutations. This may be due to the masking of recessive deleterious effects, increasing the overall performance of a variety, while maintaining recessive genetic load. One confounding factor in this conclusion could be the overall increased heterozygosity present among cultivated beet accessions compared to wild accessions (Fig. S5). The reference genome being derived from a sugar beet accession may impose some reference bias; however, the fewer mutations found in the wild ancestor accessions would suggest this would not be the primary reason for fewer deleterious mutations in the sugar beet accessions.

Although the evidence indicates that domestication of beets has increased genetic load, our results demonstrate that generations of selection and breeding have been effective at purging deleterious mutations. Sugar beet accounts for the largest share of land use, financial investment, and breeding efforts devoted to beet production relative to other cultivated beet crops (Goldman and Janick 2021). This sustained focus on sugar beet production (and improvement through breeding) over the last 1–2 centuries may explain why we detected significantly fewer deleterious mutations in sugar beet compared with the other crop types (Fig. 3B). This comparison could imply that the bottleneck resulting from selection on sugar content in the last 1700s was not of the scale or quality to increase overall genetic load as seen from the original domestication. The continued breeding and selection pressure in sugar beet is also measurable as we observed a significant decrease in deleterious mutation abundance over the past 100+ years of NPGS sugar beet entries (Fig. 4). Overall, our results indicate that these deleterious mutations may be under strong selection pressure in sugar beet breeding.

Understanding the effect of the measured deleterious mutations in modern sugar beet varieties may give some insight into variety improvement. In the subset of sugar beet lines with detailed phenotypic traits, correlations with genetic load suggest a physiological impact of these mutations (Fig. 5). The significant negative correlation between biomass and homozygous load (p<0.005) suggests the recessive action of the negative mutations are impacting overall plant fitness and has been observed before in cassava (Long *et al*. 2023). These correlations are stronger than that of measuring inbreeding alone, which implies they may be corresponding to the underlying mutations responsible for inbreeding depression. The positive relationship between sugar content and genetic load could be the product of beneficial mutations selected under domestication (Dwivedi *et al*. 2023) or because of the known negative relationship between crop yield and sugar content (Baran and YalinkiliÇ 2025). Future studies may focus on the possible subset of deleterious mutations corresponding to either original domestication or later selection on sugar content.

## Conclusion

By integrating whole‐genome sequence data from multiple studies, we were able to analyze genetic variation across almost two-thousand beet accessions. We utilized three different scales of evolutionary signal, each with their own strengths, to identify a conservative estimate of deleterious variability across *Beta vulgaris*. Our analyses reveal both the historical accumulation of deleterious mutations associated with domestication and the subsequent purging of harmful alleles through sustained sugar beet improvement efforts. The observed associations between genetic load and key agronomic traits underscore the biological relevance of these variants and highlight their potential role in inbreeding depression and performance variation. Together, these results demonstrate the value of using deep evolutionary modeling to better understand the genomic consequences of mutation. As predictive methodologies continue to advance, similar approaches will further refine our understanding of deleterious variation and support more efficient, genomics‐informed breeding and genetic engineering strategies in beets improvement.

## Data Availability

All sequence data for this project was downloaded from previously produced datasets that can be found in Table S1. Intermediate files including variant call format (vcf) file of coding genotypes along with evolutionary conservation rates, SIFT predictions, PlantCAD2 predictions, and allele frequencies can be found on the Zenodo repository: 10.5281/zenodo.19076305. For access to the larger genomic vcf (∼1Tb, 38M variants) for related research purposes, please reach out to the corresponding author.

## Acknowledgements

The authors would like to thank the data made available by the authors of the original genotyping studies. We also thank the NCBI database resource, especially with the utility of the NCBI *datasets* tools.

